# Evidence of hybridization, mitochondrial introgression and biparental inheritance of the kDNA minicircles in *Trypanosoma cruzi I*

**DOI:** 10.1101/767368

**Authors:** Fanny Rusman, Noelia Floridia-Yapur, Paula G. Ragone, Patricio Diosque, Nicolás Tomasini

**Affiliations:** Unidad de Epidemiología Molecular (UEM), Instituto de Patología Experimental, Universidad Nacional de Salta-CONICET, Salta, Salta, Argentina

## Abstract

**Background:** Genetic Exchange in *Trypanosoma cruzi* is controversial not only in relation to its frequency but also in relation to its mechanism. A mechanism of parasexuality has been proposed based on laboratory hybrids, but population genomics strongly suggests meiosis. In addition, mitochondrial introgression has been reported several times in natural isolates although its mechanism is not clear. Moreover, hybrid DTUs (TcV and TcVI) have inherited at least part of the kinetoplastic DNA (kDNA = mitochondrial DNA) from both parents.

**Methodology/Principal findings:** In order to address such topics, we sequenced and analyzed fourteen nuclear DNA fragments and three kDNA maxicircle genes in three TcI stocks which are natural clones potentially involved in events of genetic exchange. We also deep-sequenced (a total of 6,146,686 paired-end reads) the hypervariable region of kDNA minicircles (mHVR) in such three strains. In addition, we analyzed the DNA content by flow cytometry to address cell ploidy. We observed that most polymorphic sites in nuclear loci showed a hybrid pattern in one cloned strain and the other two cloned strains were compatible as parental strains (or nearly related to the true parents). The three clones have almost the same ploidy and the DNA content was similar to the reference strain Sylvio (an almost diploid strain). Despite maxicircle genes evolve faster than nuclear housekeeping ones, we did not detect polymorphism in the sequence of three maxicircle genes showing mito-nuclear discordance. In addition, the hybrid stock shared 66% of its mHVR clusters with one putative parental and 47% with the another one. In contrast, the putative parental stocks shared less than 30% of the mHVR clusters among them.

**Conclusions/significance:** The results suggest a reductive division, a natural hybridization, biparental inheritance of the minicircles in the hybrid and maxicircle introgression. The models including such phenomena and that would explain the relationships between these three clones are discussed.

**Author summary:** Chagas disease, an important public health problem in Latin America, is caused by the parasite *Trypanosoma cruzi*. Despite it is a widely studied parasite, several questions about the biology of genetic exchange remain. Meiosis has not been yet observed in laboratory, although inferred from population genomic studies. In addition, previous results suggest that the mitochondrial DNA (called kDNA) may be inherited from both parents in hybrids. Here, we analyzed a hybrid strain and the potential parents to address about the mechanisms of genetic exchange at nuclear and mitochondrial level. We observed that the hybrid strain has heterozygous patterns and DNA content compatible with an event of meiosis. In addition, we observed that the evolutionary histories of nuclear DNA and maxicircles (a part of the kDNA) were discordant and the three strains share identical DNA sequences. Mitochondrial introgression of maxicircle DNA from one genotype to another may explain this observation. In addition, we detected that the hybrid strain shared minicircles (another part of the kDNA) with both parental strains. Our results suggest that hybridization implied meiosis and biparental inheritance of the kDNA. Further research is required to address such phenomena in detail.

## Introduction

The trypanosomatids cause devastating diseases around the world [1-3]. *Trypanosoma cruzi, Trypanosoma brucei* and *Leishmania* species are the main human pathogens among the trypanosomatids, although other members of this family are also relevant [4, 5]. Genetic exchange has been proposed naturally occurring in these species [6], although its frequency is still debated [7-16]. Genetic exchange by meiosis has been described in *T. brucei* [17, 18] and *Leishmania* [19]. In spite of being recently proposed by genomic studies [14, 20] meiosis has not been reported for *T. cruzi* in the laboratory yet. Alternatively, experimental tetraploid or sub-tetraploid hybrids were observed in laboratory conditions [21, 22]. The fusion of diploid cells to form a tetraploid and posterior random loss of chromosomes slowly returning to diploidy was proposed as a model to explain genetic exchange in this species [21]. This proposed mechanism is similar to parasexuality in *Candida albicans* [23]. Interestingly, it is expected that parasexual hybrids have several chromosomes (it is expected that one-third) in a homozygous state due to random loss of both chromosomes of the same parent. In *T. cruzi*, with more than 40 pairs of chromosomes [24], the presence of chromosomes in a homozygous state in hybrid strains it would be evident if parasexuality were the mechanism of genetic exchange. In *T. cruzi*, six major lineages have been proposed (named TcI to TcVI) [25, 26]. Particularly, TcV and TcVI are naturally occurring hybrids but they did not fit the expectancy of the parasexual model. Curiously, they appear to be heterozygous in most loci [27, 28], which fit better the hypothesis of a reductive division that occurred previous to cell fusion. A recent study based on sequencing of 45 TcI genomes from a restricted geographical area showed that both mechanisms (sexual and parasexual) may occur in natural populations [20].

Inheritance of the kDNA in hybrid *T. cruzi* strains remains as an open question. Uniparental inheritance of maxicircles in natural occurring hybrids [29] and in laboratory-made hybrids was previously suggested [21]. The hybrid DTUs TcV and TcVI share nuclear genomes with the parental DTUs TcII and TcIII; instead, maxicircle sequences in both hybrid DTUs are only derived from TcIII [30]. Instead, minicircle inheritance was difficult to address in the past because technical limitations caused by the great sequence variability of such DNAs [31]. Recently, we studied the hypervariable region of the minicircles (mHVR) by deep sequencing and observed that TcV and TcVI share minicircles with both parental DTUs (TcII as TcIII) and not just with TcIII, as observed in maxicircles. Based on these results we proposed that minicircle inheritance may be biparental in natural occurring hybrids [32]. Biparental inheritance of the kDNA has also been proposed for *T. brucei* [33-35].

Another question related to the mechanisms of genetic exchange in *T. cruzi* is the mito-nuclear discordance observed in some DTUs. This phenomenon can be described as a significant difference in levels of differentiation between nuclear and mitochondrial markers, where nuclear DNA is more structured than mitochondrial DNA (kDNA in this case) [36]. In *T. cruzi*, mito-nuclear discordance is clearly observed in TcIII. According to nuclear loci, TcIII is nearly related to TcI; although maxicircle sequences cluster TcIII with TcIV from South America [30, 37]. In addition, mito-nuclear discordance was also observed within TcI [38-40]. A probable cause of mito-nuclear discordance in *T. cruzi* is mitochondrial introgression i.e. successive backcrosses of a hybrid with one of the parents.

In previous studies, we described a TcI strain (TEDa2cl4) isolated from Chaco Province in Argentine that it is a potential hybrid. TEDa2cl4 presented some heterozygous patterns in isoenzymatic loci [41], a high number of heterozygous SNPs in the spliced-leader intergenic region [42] and heterozygous patterns in nuclear loci revealed by MLST [28, 43]. We isolated — within the same restricted geographical area in Chaco (Argentina)— two strains (TEV55cl1 and PalDa20cl3) that are potential parents considering isoenzymatic loci [41], the SL-IR [42] and MLST [28, 43]. This strain triplet is an interesting opportunity to address the mechanism of genetic exchange in *T. cruzi*. Here, we analyzed the DNA sequence for 14 nuclear loci, 3 maxicircle regions and around 6.1 million paired-end reads of the mHVRs for this triplet of TcI strains. We also addressed ploidy of the strains by flow cytometry.

## Materials and methods

### Strains

Three laboratory cloned stocks isolated from Chaco province, Argentina, and belonging to TcI, were analyzed. The stocks characteristics are summarized in Table 1. In addition, the TcI strains Sylvio and TEV91cl5 were also cultured and used as controls. The stocks were maintained in liver infusion-tryptose (LIT) medium supplemented with 10% fetal bovine serum. Parasites in exponential growth phase were harvested by centrifugation (800 xg, 10 min, 4 °C) for DNA extraction and flow cytometry. DNA was extracted by using a commercial kit (Inbio Highway).

**Table 1.** TcI stocks used in this study to address hybridization and kDNA inheritance

| Stock | Host | Geographical origin<br>(Chaco Province,<br>Argentina) | SL-IR group<sup>1</sup> / MLST cluster<sup>2</sup> |
| --- | --- | --- | --- |
| PalDa20cl3 | *Didelphis albiventris* | EL PALMAR, 12 de Octubre | Chaco-3 / Chaco-3 |
| TEDa2cl4 | *Didelphis albiventris* | TRES ESTACAS, Chacabuco | Chaco-3 / potential hybrid<sup>3</sup> |
| TEV55cl1 | *Triatoma infestans* | TRES ESTACAS, Chacabuco | Chaco-2 /Chaco-2 |

1. Group according to Spliced Leader Intergenic Region (SL-IR) sequence [42]. 2. Cluster according sequence of 8 nuclear gene fragments as analyzed in [43]. 3. MLST analysis suggested TEDa2cl4 is a hybrid between Chaco-2 and Chaco-3 Clusters [43].

### Sequencing of nuclear and maxicircle gene fragments

The primers used to amplify eight nuclear regions and three maxicircle gene fragments are shown in Table 2. PCRs were carried out in reaction volumes of 50 µl containing 50 ng of DNA; 0.2 µM of each primer, 1 U of goTaq DNA polymerase (Promega), 10 µl of 5X buffer (supplied with the goTaq polymerase) and a 50 µM concentration of each dNTP (Promega). Cycling conditions were as follow: 3 min at 94 °C followed by 30 cycles of 94 °C 30 seconds; 50 °C 30 seconds, and 72 °C for 30 seconds, with a final extension at 72 °C for 10 min. Amplified fragments were precipitated with 70% ethanol and sequenced on both strands in an ABI PRISM_310 Genetic Analyzer (Applied Biosystems). In addition, sequences from genes leucine aminopeptidase (*LAP*), glucose-6-phosphate isomerase (*GPI*), glutathione peroxidase (*GPX*), pyruvate dehydrogenase E1 component alpha subunit (PDH), 3-hydroxy-3-methylglutaryl-CoA reductase (HMCOAR) and small GTP-binding protein rab7 (GTP) were obtained in a previous work [43].

**Table 2.**
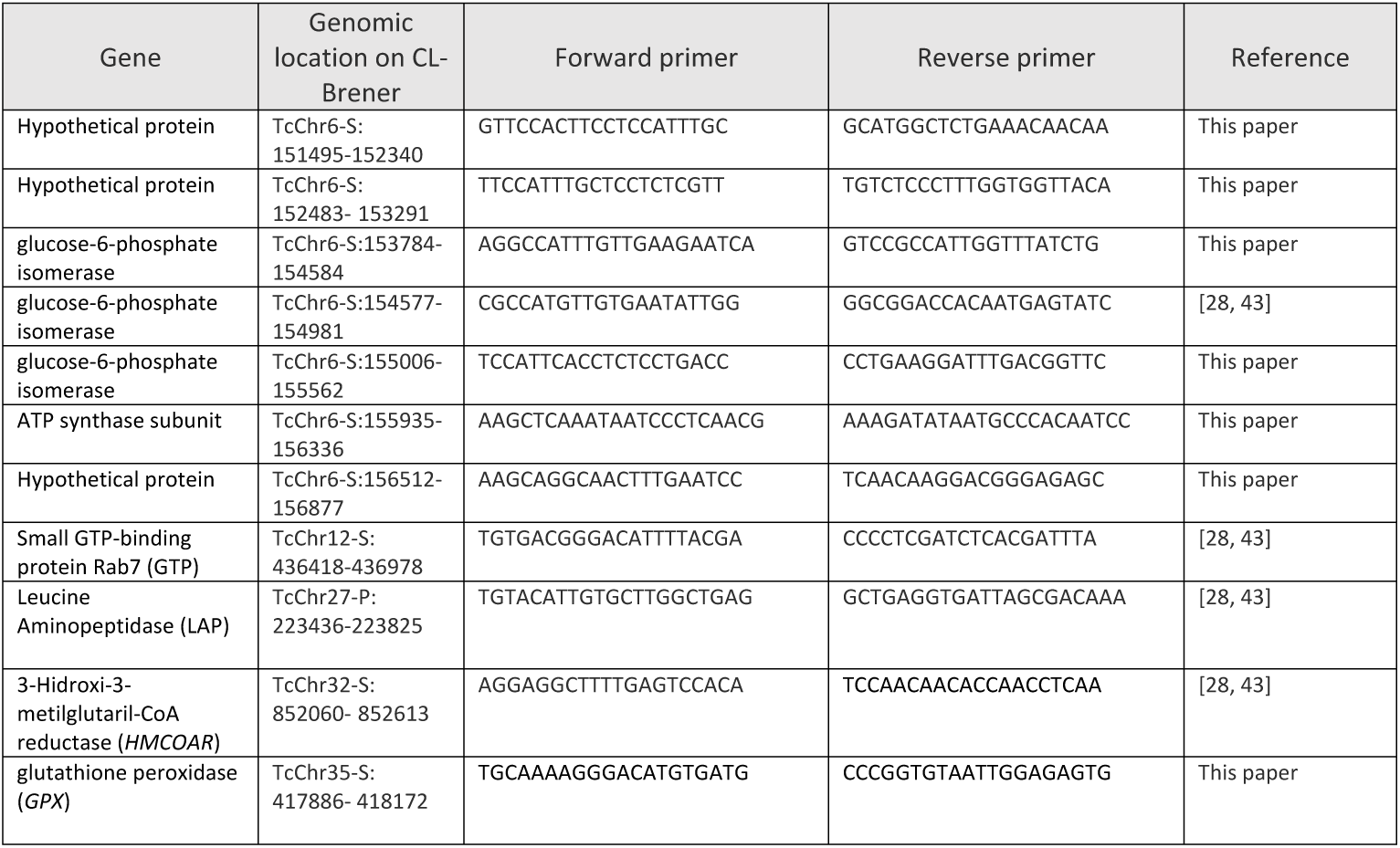
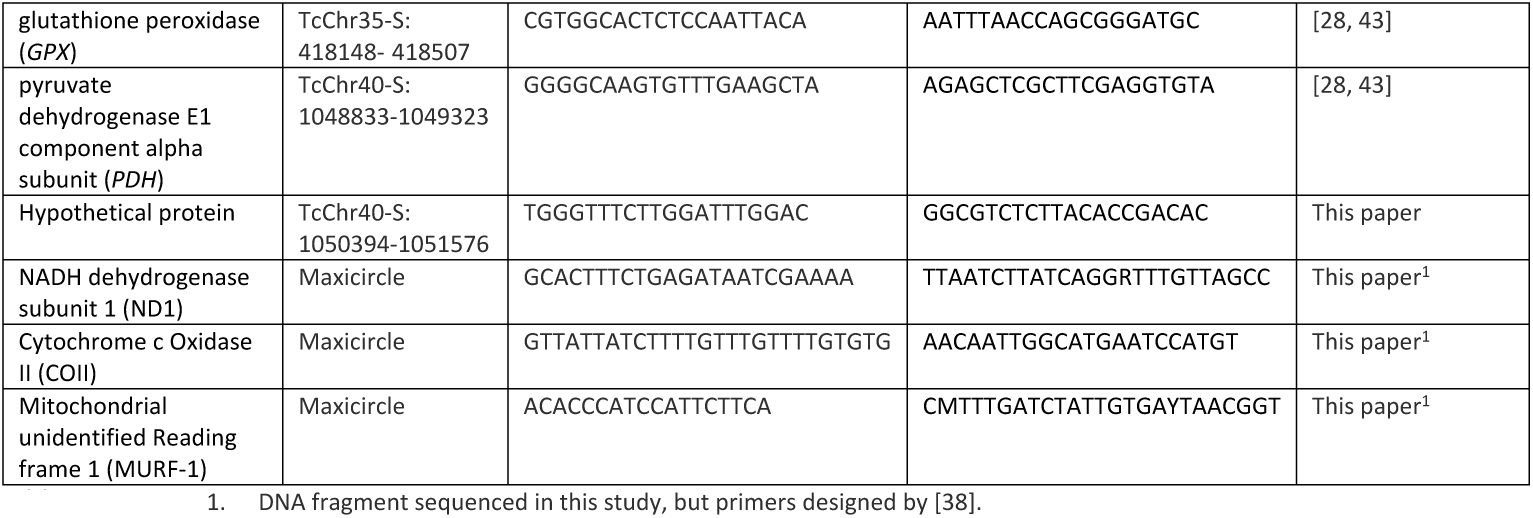
Gene fragments, genomic location and primers used in this study

### Flow cytometry

Pellets containing 1×10^7^ parasites were resuspended in 300 µl of PBS and 700 µl of ice-cold methanol. The tube was gentle inverted and incubated on ice for 10 min. Fixed parasites were centrifuged at 800 xg and washed in 5 ml of PBS two times. The pellet was resuspended in 1 ml of PBS. Propidium Iodide and RNAse A were added to a final concentration of 30 µl/ml and 10 µg/ml, respectively, and incubated 45 min at 37 °C. Samples were analyzed in a FACS Canto II in FL2 channel. Around 50,000 events were recorded for each sample and the samples were analyzed by duplicate. Flowing software (http://flowingsoftware.btk.fi) was used for data analysis. After gating out debris and cell clumps the data were plotted as fluorescence area histograms. Gates were created for G1/0 (2*n*) peaks and for G2/M (4*n*) peaks. Mean G1/0 values were used to infer relative DNA content. The coefficient of variation (CV) was recorded for each fluorescence peak. The DNA content for each stock was expressed relative to Sylvio strain, an almost diploid strain [44].

### mHVR sequencing

The mHVRs for TEDa2cl4 were amplified according the protocol and primers proposed in [32]. Amplicons were then purified using the magnetic beads Agencourt AMPure XP-PCR Purification (Beckman Genomics, USA). The concentration of the purified amplicons was controlled using Qubit Fluorometer 2.0 (Invitrogen, USA). The library was validated using the Fragment Analyzer system (Advanced Analytical Technologies, USA). The library was sequenced on an Illumina MiSeq using a 500 cycle v2 kit (Illumina, San Diego, USA). The mHVR reads from TEV55cl1 and PalDa20cl3 were previously published in [32].

### Bioinformatics

The reads were demultiplexed, trimmed and filtered according to [32]. In addition, preprocessed sequences were clustered with the “pick_de_novo_otus.py” script in QIIME V1.9.1 [45]. The *de novo* approach groups sequences based on sequence identity using the uclust algorithm [46]. Default parameters were used, and sequences were clustered according to two different identity thresholds —85% and 99%— in order to determine different mHVR clusters. Output tables were filtered at 0.005% using the “filter_otus_from_otu_table.py” script, in order to discard mHVR clusters with low abundance which are probably sequencing artifacts [47], remaining parameters were used by default. The presence of a cluster into a strain was discarded when its abundance was lower than 20 read sequences. In order to compare stocks with different number of reads, the datasets were rarefacted to the number of reads in the smallest dataset (TEDa2cl4 in this case).

### Phylogenetic analysis

In order to address mito-nuclear discordance, the uncorrected *p*-distance and a model corrected distance were calculated between pairs of strains (TEV55cl1-CL Brener, TEV55c1-Esmeraldo, CL Brener-Esmeraldo, TEV55cl1-TEV91cl5 and TEV55cl1-PalDa20cl3) for each gene fragment and by using MEGA v7 [48]. The CL Brener sequences for each nuclear gene fragment were obtained from CL Brener (non Esmeraldo) genome [24] from TriTryp. The average distance (model corrected and uncorrected) and the standard deviation were calculated for nuclear and maxicircle DNA fragments in order to compare substitution rates. Model corrected distances were calculated using MEGA v7 software by selecting the best substitution model for each gene fragment and calculating the pairwise distance between strains according to the selected model. In addition, sequences for the 14 nuclear gene fragments and three maxicircle genes from TcI strains JRcl4 and Dm28c (available genomes in TriTryp) were included into the analyses in order to address mito-nuclear discordance in TcI. Nuclear and maxicircle sequences were concatenated in two independent alignments using MLSTest [49]. The concatenated sequences were analyzed using MEGA v7 to select the best substitution model and to build phylogenetic trees with 1,000 bootstrap replications.

### Accession Numbers

Nuclear DNA sequences from [43] were downloaded from GenBank (KF264037, KF268604, KF268636, KF268620, KF264005, KF264021, JN129573, JN129606, JN129639, JN129507, JN129540, JN129672, JN129608, JN129509, JN129674, JN129641, JN129575, JN129542). Nuclear and maxicircle DNA sequences obtained in this paper are available in GenBank with the following accession numbers: (MN413539-MN413581). The mHVR reads for the three strains are available the NCBI SRA database (BioProject ID: PRJNA514922).

## Results

### Analysis of fourteen nuclear loci and DNA content reveals that TEDa2cl4 is an almost diploid hybrid clone

A total of 7,135 bases corresponding to 14 gene fragments on 6 chromosomes were analyzed in the stocks TEDa2cl4, PalDa20cl3 and TEV55cl1. The polymorphic sites for each fragment are shown on Fig 1A. The number of polymorphisms was low (36 — 0.5% of the total analyzed sites). Thirty (83.3%) polymorphic sites were heterozygous in TEDa2cl4. Moreover, in 29/30 (96.7%) of these heterozygous sites, the pattern is compatible with the hypothesis of PalDa20cl3 and TEV55cl1 as parents (or at least closely related to the parents). This result reinforces the hypothesis of TEDa2cl4 as a hybrid. Additionally, heterozygous SNPs in TEDa2cl4 were found in different chromosomes (Fig 1A). Instead, the percentage of heterozygous polymorphic sites was relatively lower in PalDa20cl3 (25%) and TEV55cl1 (13.9%) than in TEDa2cl4. The flow cytometry analysis for the three strains showed that they have similar DNA contents (Fig 1B) suggesting a reductive division as meiosis. Particularly, the DNA content of TEDa2cl4 (hybrid) and TEV55cl1 (putative parental) were almost identical. The estimated ploidy level is similar to the obtained in the almost diploid reference strain Sylvio (Fig 1C). In addition, PalDa20cl3 showed the higher relative DNA content (2.3n expressed as the relative DNA content in relation to a half of the DNA content on Sylvio).

**Fig 1.**
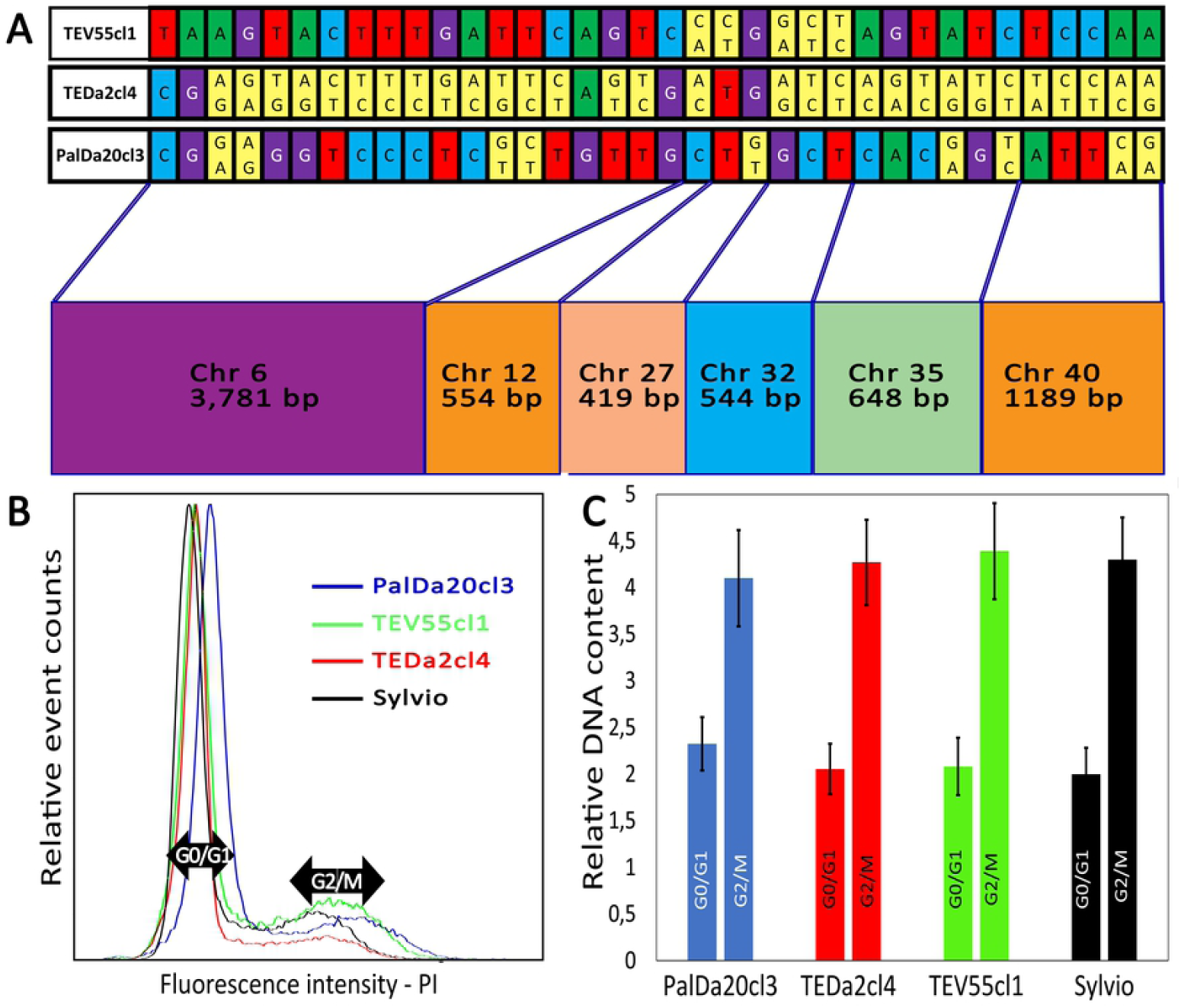
TEDa2cl4 is an almost diploid hybrid strain. (A) polymorphic sites in 14 nuclear DNA fragments for the strains TEV55cl1, TEDa2cl4 and PalDa20cl3 showing heterozygous sites (yellow bases). (B) Proportion of cells with different DNA content measured by fluorescence intensity of propidium iodide in flow cytometry. (C) DNA content in different cell-cycle stages in relation to the almost diploid strain Sylvio.

### Maxicircle genes analysis showed mito-nuclear discordance

In contrast to the polymorphic sites observed in the nuclear loci, sequences of three maxicircle genes (1,335 bp) revealed no polymorphic sites among the three strains. This was an unexpected result because coding maxicircle genes mutate faster than nuclear coding genes. This is revealed in Fig 2A showing such different substitution rates among different DTUs and within TcI. When uncorrected *p*-distance or model corrected distances were considered for pairs of strains, a higher average substitution rate in maxicircle loci than in nuclear loci was observed for most comparisons (Fig 2A and Fig S1). The exception was the pair TEV55cl1-PalDa20cl3, which had more substitutions at nuclear loci than at mitochondrial loci. This result indicates mito-nuclear discordance. A comparison between nuclear and mitochondrial phylogenetic trees also reveals the mito-nuclear discordance (Fig 2B). In addition, the trees also showed discordance considering other TcI strains suggesting that it is not an uncommon phenomenon.

**Fig 2.**
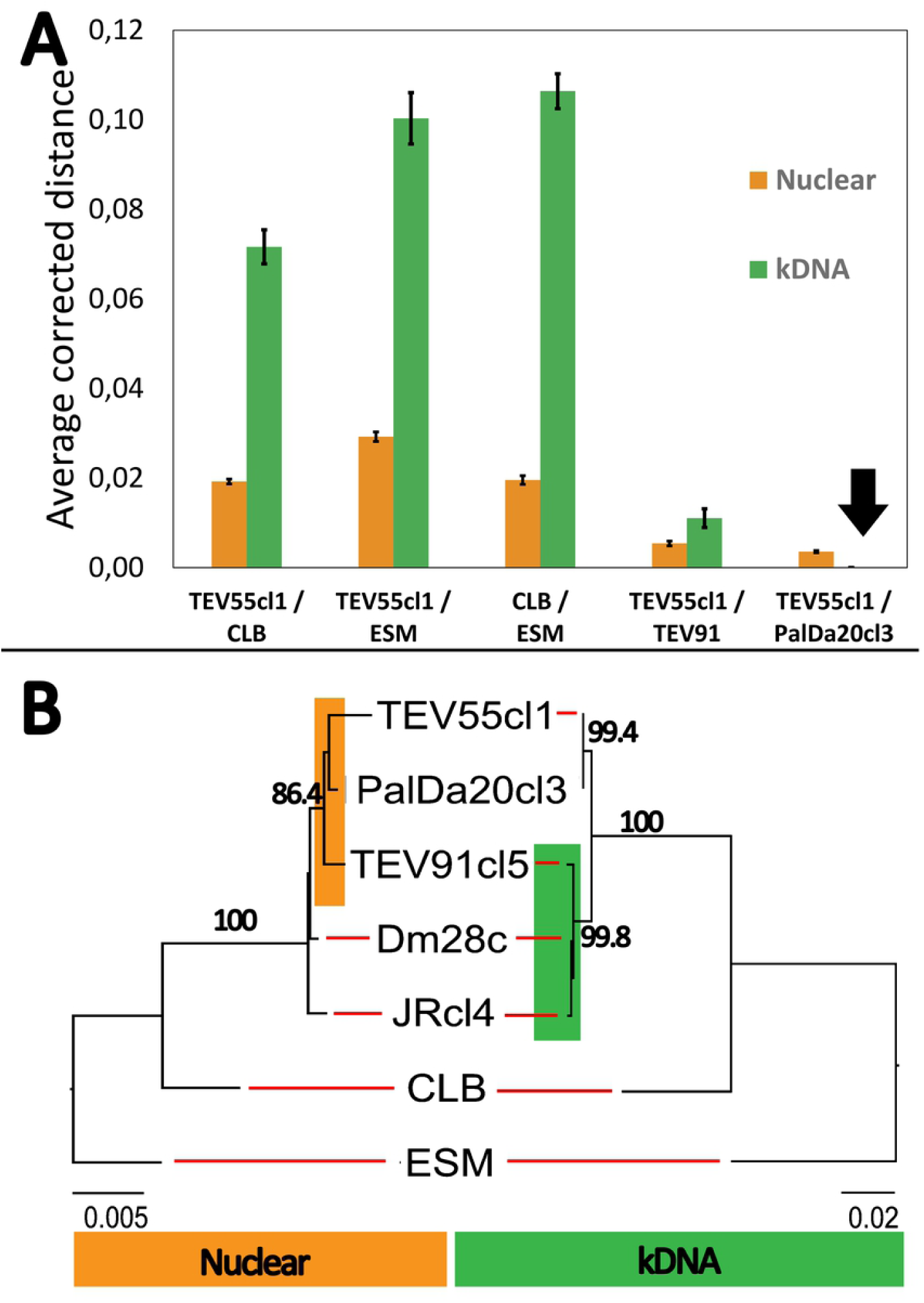
Mito-nuclear discordance in TcI strains. (A) Comparison of averaged model-corrected distances among nuclear loci and kDNA loci. Sequences from TcI (TEV91cl5, TEV55cl1, PalDa20cl3), TcII (Esmeraldo-ESM), and TcIII (CLB, inferred from CL-Brener Non Esmeraldo genome) where used in order to address relative substitution rates. Most comparisons showed higher genetic distances for kDNA markers than for nuclear loci except for the comparison between TEV55cl1 and PalDa20cl3. Error bars represent the standard error of the mean corrected distance. (B) Nuclear phylogeny (left) and kDNA phylogeny (right) showing discordances between both markers. The trees were estimated based on the best substitution model for concatenated genes. Values above branches indicate bootstrap support for 1,000 replicates. Note that location in different clusters of TEV91cl5 (highlighted) and different branch scales for both markers.

### mHVR deep sequencing reveals that minicircles are biparentally inherited in TEDa2cl4

We performed deep sequencing of the mHVR amplicons for the three strains. The number of reads for each strain is observed in Table 3. Sequences were clustered according the percentage of sequence identity. Despite these strains had no variation in the three maxicircle loci analyzed (Fig 2), the number of mHVR clusters was highly variable among strains (Table 3). In addition, a rarefaction analysis showed that the number of clusters did not depend on the number of reads for each strain.

**Table 3.** Deep sequencing of the minicircle hypervariable region (mHVR)

|  | TEV55cl1 | TEDa2cl4 | PalDa20cl3 |
| --- | --- | --- | --- |
| Number of paired Reads | 2,305,460 | 1,113,033 | 2,728,193 |
| Filtered and merged | 1,202,355 | 950,142 | 2,356,494 |
| Number of mHVR clusters | 285 | 217 | 350 |
| Number of mHVR clusters rarefacted to 900,000 reads (standard deviation for 100 replicates) | 284.8(0.4) | 215.6(0.9) | 346.8(0.4) |

A Principal Coordinates Analysis (PCoA) (Fig 3A-B) of Bray-Curtis dissimilarities among strains revealed that TEDa2cl4 did not cluster closely to one of the parental-like strains as would be expected under the hypothesis of uniparental inheritance of minicircles. Conversely, PCoA showed a TEDa2cl4 located almost at the middle between both strains (Fig 3B). In addition, we analyzed shared clusters at different identity thresholds (Fig 3C-D). At 85% identity threshold, most of the mHVR clusters in TEDa2cl4 (84%) were observed in TEV55cl1 and/or PalDa20cl3 (Fig 3C) suggesting biparental inheritance of the minicircles in this hybrid. Despite, TEV55cl1 and PalDa20cl3 had higher percentages (40.4% and 63.1% respectively) of mHVR clusters which are singletons (i.e. they are not shared with any other strain) (Fig 3C). We also analyzed shared mHVR clusters at a more restrictive clustering threshold (at 99% identity threshold). In this case, the number of mHVR clusters in TEDa2cl4 that were observed in PalDa20cl3 and/or TEV55cl1 was reduced to 50% (Fig 3D). This genetic divergence suggests that it is more likely that TEV55cl1 and PalDa20cl3 are not related to TEDa2cl4 in a recent time (i.e. they are not direct parents). However, at this threshold, it is interesting that TEV55cl1 and PalDa20cl3 almost did not share any mHVR cluster (8 mHVR clusters which correspond to 2.9% and 2.2% respectively), although they still share a significant percentage of mHVR clusters with the hybrid TEDa2cl4 (Fig 3D). Interestingly, from the mHVR clusters that TEDa2cl4 shared with any other strain, 54% were observed in TEV55cl1 but not in PalDa20cl3 and 42% were observed PalDa20cl3 but not in TEV55cl1. Such results strongly support that minicircle inheritance is biparental in the hybrid strain.

**Fig 3.**
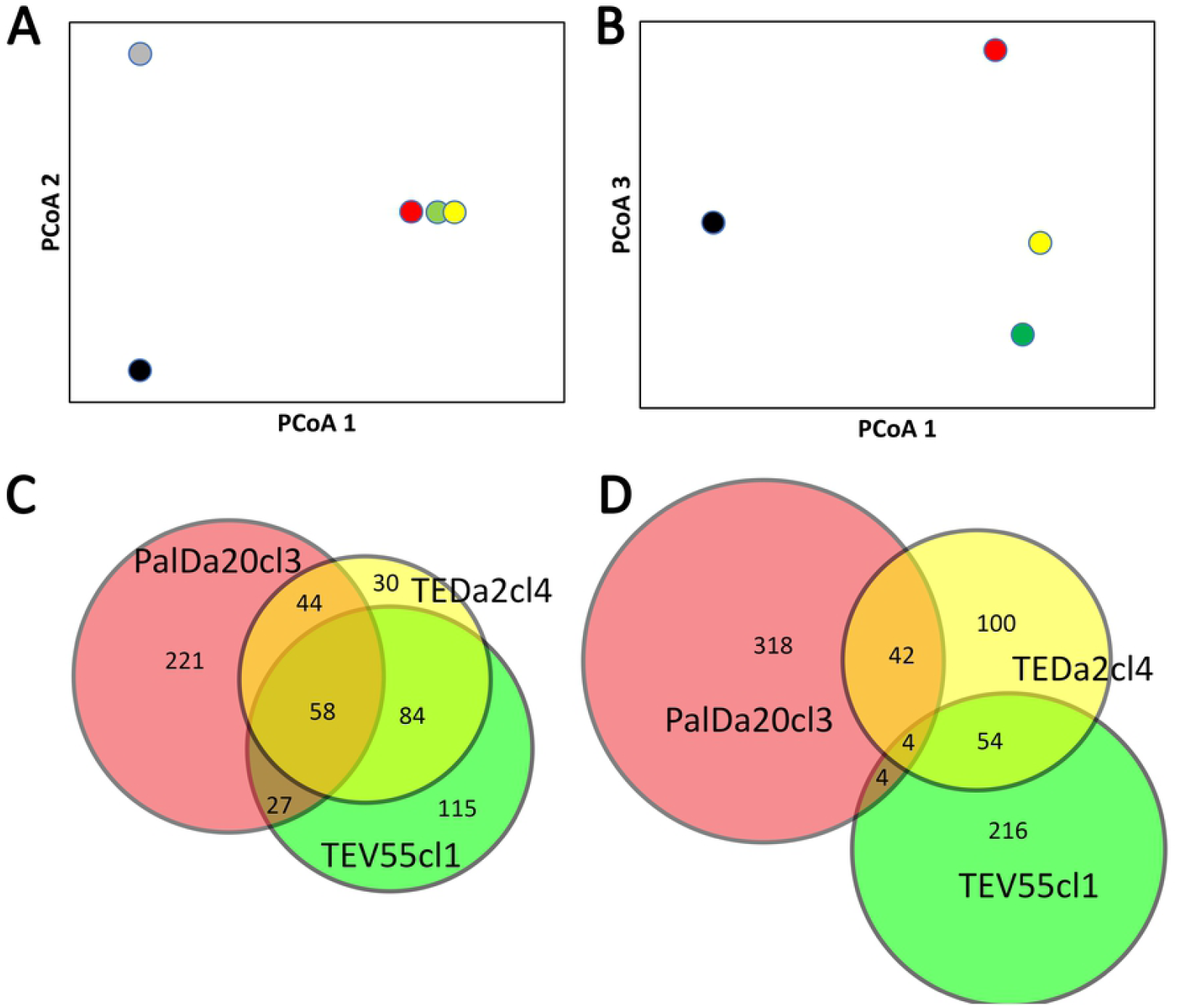
Minicircles of the hybrid strain TEDa2cl4 are biparentally inherited. (A-B) Principal Coordinates Analysis based on Bray-Curtis dissimilarities for clusters at 85% identity threshold showing the first three axes for PalDa20cl3 (red dot), TEV55cl1 (green dot), TEDa2cl4 (yellow dot) and the outgroup strains CANIIIcl1 (TcIV, grey dot) and X109/2 (TcIII, black dot). The three first axes explain 84% of the observed dissimilarities. (A) Axis 1 vs Axis 2, the first two axes show the separation among different DTUs. (B) the third axis show the separation between TcI strains. Note that TEDa2cl4 is located between PalDa20cl3 and TEV55cl1. (C-D) Analysis of shared mHVR clusters between the thee strains using Venn diagrams. Each circle represents the total number of mHVR clusters in a strain. Overlapped regions among circles represent shared mHVR clusters. (C) mHVR clusters defined at 85% of identity threshold showing few singletons in TEDa2cl4. (D) mHVR clusters defined at 99% identity threshold.

## Discussion

The genetic exchange has been reported in natural populations of *T. cruzi* [14, 20, 38, 39, 50-52]. Meiosis has been proposed in genomic population studies [14, 20] although the experimental observation of gametes and laboratory meiotic hybrids is still elusive. In addition, the frequency of genetic exchange and how the population genetic structure is maintained is still under debate [7-16, 20]. Here, we analyzed three strains that previously showed evidence related to genetic exchange events [28, 41, 42, 53]. In these three strains we analyzed DNA sequences at nuclear, maxicircle and minicircle levels. Interestingly, the results showed that the strain TEDa2cl4 is the result of a hybridization between different TcI genotypes and it maintains diploidy suggesting a reductive division. The strains TEV55cl1 and PalDa20cl3 are, at least, genetically related to the parental genotypes of TEDa2cl4. In addition, analysis of maxicircle genes reveals mito-nuclear discordance and suggests mitochondrial introgression by backcrosses in TEV55cl1 and/or PalDa20cl3. Finally, we observed that the hybrid TEDa2cl4 share minicircles (mHVRs) with both putative parental genotypes, which strongly suggests biparental inheritance of minicircles in hybrids.

The analysis of nuclear DNA fragments of at least five different chromosomes revealed that TEDa2cl4 is highly heterozygous suggesting that it was maintained by clonal propagation. We have also detected other strains with the same genotype than TEDa2cl4 in the same area [53]. A recent paper has revealed the existence of nuclear genetic exchange in natural populations of TcI in Arequipa, Peru [14]. The authors proposed inbreeding as the main cause for determining the genetic structure of this lineage. However, the TcI genomes from Arequipa in the above-mentioned paper showed high levels of heterozygosity, an improbable feature in populations with high levels of inbreeding [54]. Our data also do not support the hypothesis of inbreeding as the main determinant of the genetic structure because of the highly heterozygous nature of TEDa2cl4 and related strains. Moreover, our previous results for TcI in a restricted geographical area revealed well defined clusters with high linkage disequilibrium, congruence among different genetic markers and no geographical association [53]. However, another population genomic study in TcI stocks from Ecuador also showed that almost panmictic populations and clonal populations may coexist [20].

The mechanism of genetic exchange is still debated. The model of parasexuality proposes diploid fusion and posterior chromosome loss to return diploidy [21]. Fusion of diploid cells has been observed in the laboratory [21]. However, the returning from tetraploidy to diploidy was never observed in such hybrids [55]. Parasexuality appears not to be frequent in nature although it cannot be discarded from data of a population genomic study where several aneuploid stock were observed [20]. Despite, hybrid DTUs such as TcV and TcVI have heterozygous patterns more compatible with meiosis and gamete fusion than with the model of parasexuality [37, 55]. An expected result of the parasexual model is the existence of strains with high degrees of aneuploidy. Contrary, our results suggest meiosis since the hybrid TEDa2cl4 has a DNA content compatible with diploidy: TEDa2cl4 has the same DNA content than an almost diploid strain (Sylvio has 1/41 pairs of chromosomes in aneuploidy) [44]. Despite, sexual and parasexual modes of reproduction may be occurring simultaneously in natural populations [20].

The three strains also give us the possibility to address kDNA inheritance in hybrids. Maxicircle inheritance is assumed to be uniparental according to previous results in hybrid DTUs TcV and TcVI, since they share maxicircles with TcIII but not with TcII [30, 37]. Our data did not allow to address the hypothesis of uniparental inheritance of maxicircles because the DNA fragments analyzed here were identical among the three strains. This was surprising since substitution rates are higher in kDNA coding genes than in nuclear ones. Under neutral theory, the changes in nuclear and maxicircle DNAs are accumulated with different speed, but the ratio between the magnitude of accumulation is expected to be the same regardless of divergence time between the lineages [36]. This is not the case here. Theoretically, maxicircles could have become fixed in the ancestral strain and retained unchanged in the diverged genotypes for stochastic reasons or stabilizing selection. However, this last scenario is unlikely considering the accumulated mutation rate in maxicircles (See Fig 2A). In addition, low divergency in maxicircles was also reported in TcI populations where sexual reproduction has been observed [14, 20]. Consequently, mitochondrial introgression is the most likely explanation. Mitochondrial introgression consists in successive backcrosses of the hybrid with one of the parental genotypes. The successive backcrosses clean the genome of information from one parental. However, maxicircles from such parental are maintained (see Fig 4). According to such hypothesis, TEV55cl1 and/or PalDa20cl3 are probably descendants of hybrids that backcrossed with one parental, which could explain why such strains share identical maxicircle sequences (Fig 4).

**Fig 4.**
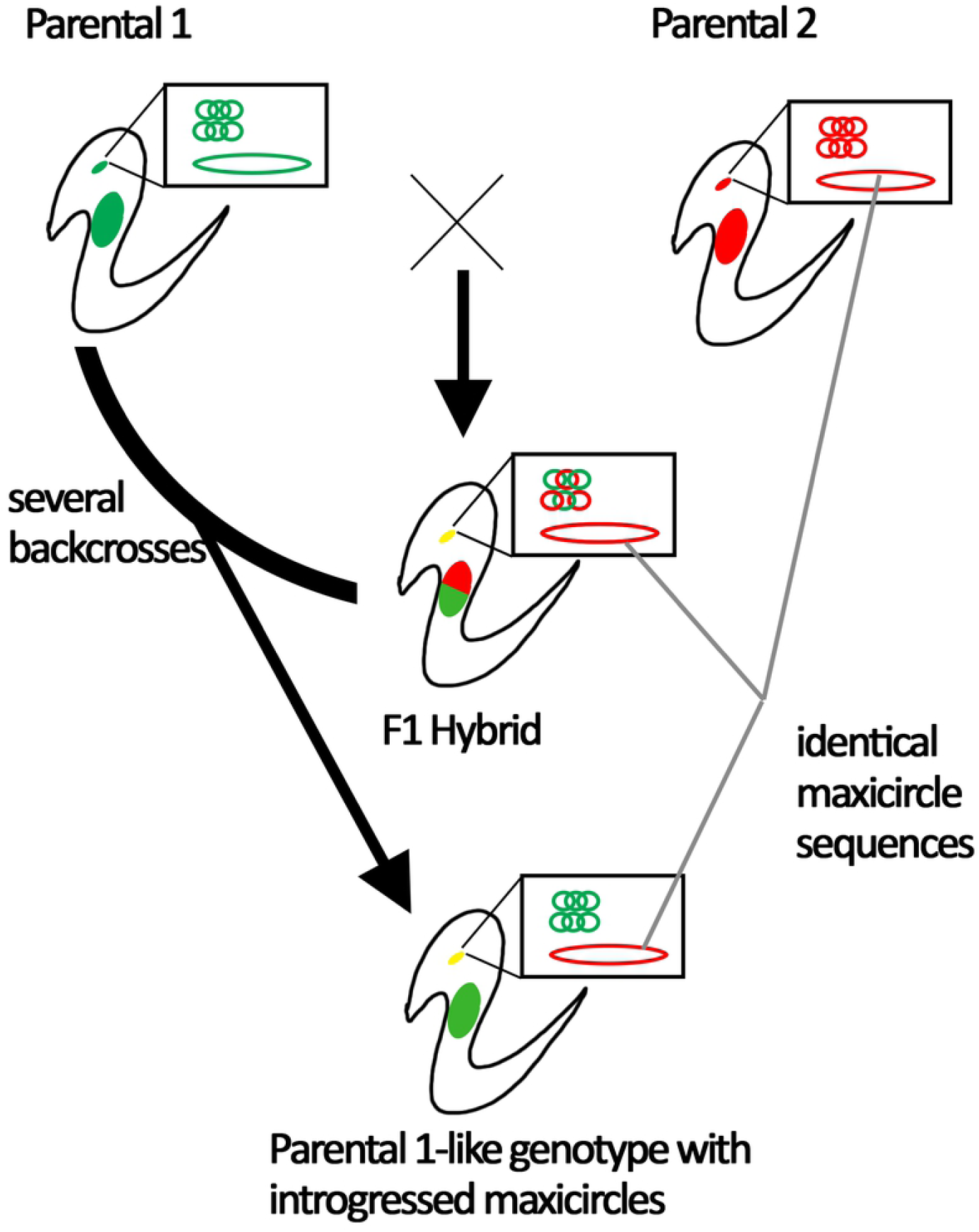
Model explaining shared maxicircles among the TEDa2cl4, TEV55cl1 and PalDa20cl3. Two parental strains with different nuclear genomes (big red and green solid ellipses), different maxicircles (big red and green unfilled ellipses) and different minicircles (catenated small circles) crosses to form an F1 hybrid. The F1 has mixes of minicircles and both nuclear genomes but maxicircles from just one parental. Backcrosses of the F1 get the descendants of the F1 more and more similar to the one of the parental genotypes but maintaining the maxicircle from the other parental.

In this work, the inheritance of minicircles was also addressed. We evaluated the number of shared mHVR clusters between strains. We observed that most of the mHVR clusters in TEDa2cl4 (84%) were shared with TEV55cl1 and/or with PalDa20cl3 at 85% identity threshold. Instead, TEV55cl1 and PalDa20cl3 shared less than 30% of the mHVR clusters. These results support that TEDa2cl4 has minicircles from two different sources corresponding to two different genotypes. Our results also show that TEDa2cl4 is not a direct descendant of PalD20cl3 and TEV55cl1 considering that there was a 50% of mHVR clusters in TEDa2cl4 that were not present in the studied putative parental strains when a restrictive identity threshold is used (i.e. 99% of sequence identity).

We have previously suggested biparental inheritance of minicircles in TcV and TcVI [32], which, added to the results presented here, suggests that biparental inheritance of minicircles could be the rule in *T. cruzi*. In *T. brucei*, it has been observed that maxicircle and minicircle inheritance is biparental in hybrids. However, maxicircles (less than fifty copies) are homogenized by genetic drift resulting in the loss of maxicircles of one parental in few generations. However, minicircles have much more copies (thousands) and they resist the fixation effect of genetic drift for more time. Consequently, in *T. brucei* maxicircle inheritance is biparental and just seems to be uniparental due to genetic drift [33-35].

Concluding, this paper shows a natural TcI hybrid and it supports that genetic exchange in *T. cruzi* likely implies a reductive division such as meiosis. In addition, genetic exchange also implies kDNA exchange. In this sense, events of mitochondrial introgression appear to be frequent in the population which is an evidence of outcrossing instead of inbreeding. This probably occurs in regions where the different genotypes overlap their distributions. The mechanism of mitochondrial DNA exchange needs to be clarified in future studies in order to resolve if fusion of mitochondria is required to biparental inheritance of minicircles. In addition, investigating on meiosis, gametes and experimental F1 hybrids is still needed.

## Supporting information

Fig S1. Comparison of averaged uncorrected *p*-distances among nuclear loci and kDNA loci.

## Acknowledgements

We would like to thank Andrea Puebla, Pablo Alfredo Vera, Veronica Nishinakamasu and Marianne Muñoz (Instituto de Biotecnología, Centro de Investigaciones en Ciencias Agronómicas y Veterinarias, Instituto Nacional de Tecnología Agropecuaria) for their technical assistance in mHVR sequencing.

